# 3D human retinal organoid model for the study of early diabetic retinopathy

**DOI:** 10.1101/2023.09.04.556258

**Authors:** Luisa de Lemos, Pedro Antas, Inês S. Ferreira, Inês Paz Santos, Catarina M. Gomes, Catarina Brito, Miguel C. Seabra, Sandra Tenreiro

## Abstract

Diabetic retinopathy (DR) is a significant complication of diabetes and a primary cause of visual impairment among working-age individuals. DR is a degenerative condition in which hyperglycaemia results in morphological and functional changes in certain retinal cells. Existing treatments mainly address the advanced stages of the disease, which involve vascular defects such as macular edema or neovascularization. However, it is now known that retinal neurodegeneration and inflammation precede these vascular changes. Therefore, there is a pressing need to identify new therapeutic approaches that target the early stages of DR and prevent its progression.

In the last decade, the development of reliable *in vitro* models resembling the complexity of the retinal tissue has significantly improved. Namely, three-dimensional (3D) retinal organoids derived from human induced-pluripotent stem cells (hiPSCs) recapitulate the cellular organization and complexity of the human retina.

Here, we used hiPSCs-derived retinal organoids to generate a model of early DR. In this model, we observe well-established molecular and cellular features of early DR: *i*) loss of retinal ganglion and amacrine cells; *ii*) glial reactivity and inflammation, with increased expression of the vascular endothelial-derived growth factor (*VEGF)* and interleukin-1β (*IL-1β),* and monocyte chemoattractant protein-1 (MCP-1) secretion; *iii*) increased levels of reactive oxygen species accompanied by activation of key enzymes involved in antioxidative stress response. The data provided highlight the utility of retinal organoid technology in modelling early-stage DR. This offers new avenues for the development of targeted therapeutic interventions on neurodegeneration and inflammation in the initial phase of DR, potentially slowing the disease’s progression.

## Introduction

Diabetic retinopathy (DR) is a common complication of diabetes and a leading cause of vision loss worldwide (Sivaprasad et al., 2012; Lee et al., 2015; Kropp et al., 2023). The traditional model of DR as a microvascular disease has evolved. It is now well-established that neurodegeneration is not only significant in DR but is also considered an early event, preceding visible microvascular abnormalities. This fact is referred to as diabetic retinal neurodegeneration (DRN) (Chakravarthy and Devanathan, 2018; Zafar et al., 2019; Soni et al., 2021). However, there are no treatments targeting the early stages of the disease before the onset of vascular defects such as macular edema or neovascularization or before retinal neurodegeneration occurs (Sohn et al., 2016; Simó et al., 2018; Channa et al., 2021). Although reactive gliosis, neural apoptosis and microvascular dysfunction can be interdependent and essential for developing DR, neuroprotection can be considered a therapeutic target per se (Chakravarthy and Devanathan, 2018; Zafar et al., 2019). Consequently, it is of paramount importance to elucidate the intricate molecular and cellular mechanisms underpinning DRN and to develop novel therapeutic approaches targeting its early stages, thereby mitigating the disease’s progression.

Hyperglycaemia, hypertension and diabetes duration are considered the established risk factors for developing DR (Brownlee, 2001; Nathan, 2014; Hammer and Busik, 2017). Several cellular pathways have been proposed to explain DR. The most studied mechanisms are increased polyol and hexosamine pathways flux, increased advanced glycation end-products (AGE) formation, abnormal activation of protein kinase C (PKC) pathway and increased oxidative stress (Brownlee, 2001). Common grounds for all these mechanisms are oxidative stress, inflammation, vascular occlusion, upregulation of factors such as insulin-like growth factor (IGF), vascular endothelial growth factor (VEGF), tumor necrosis factor (TNF) and basic fibroblast growth factor-2 (bFGF). The activation of these pathways leads to the degeneration of the neural retina and capillary abnormalities in the inner retina. Neural apoptosis and reactive gliosis are considered the most important histological features of DR. Retinal ganglion cells (RGCs), located in the inner retina, are the more susceptible cells to hyperglycaemia (Kern and Barber, 2008), and RGCs loss has been detected in diabetic rats and diabetic patients either without or with only minimal DR (Sima et al., 1992; Barber et al., 1998; Asnaghi et al., 2003; Kern and Barber, 2008; van Dijk et al., 2009, 2012). In addition to RGCs, amacrine cells and photoreceptors have showed an increased apoptotic rate in diabetic retinas (Barber et al., 1998; Carrasco et al., 2007; Garcia-Ramírez et al., 2009).

In the last decades, *in vitro* two-dimensional (2D) models of DR have contributed to characterize the cellular processes of retinal damage during diabetic conditions. These models provide simpler systems of one or two cell types in interaction for studying the cytotoxic effects of high glucose, glutamate, or AGEs (Matteucci et al., 2015). Particularly, cell cultures of dorsal root ganglion cells, cortical and hippocampal neurons, and RGCs were frequently used to highlight the early events in DR (Russell et al., 1999; Santiago et al., 2007; Fu et al., 2012). The involvement of Müller glia in DR was also explored on primary Müller cells and established Müller cell lines and have contributed to understand the proinflammatory role of Müller cells in high glucose conditions (Xi et al., 2005; Matteucci et al., 2014; Tien et al., 2017).

Organotypic retinal cultures, or retinal explants, which preserve the histotypic architecture of the retina have been established to elucidate the interplay between neuronal and glial compartment in the context of DR. Nevertheless, these models pose challenges in regard to duration and the viability of the *in vitro* culture (Caffé et al., 2002; Matteucci et al., 2015; Valdés et al., 2016).

Over the years, numerous animal models have been established to study the molecular and cellular mechanisms of DR. These include models of inducted hyperglycaemia, spontaneously diabetic rodents, genetic DR models carrying mutations in the leptin or insulin-2 genes, alongside with models of angiogenesis without diabetes (Robinson et al., 2012; Lai and Lo, 2013; Ly et al., 2014; Olivares et al., 2017). Nonetheless, the majority of these models predominantly exhibit the early characteristics of DR, while few replicate the advanced proliferative stage of DR, characterized by angiogenesis (Robinson et al., 2012; Lai and Lo, 2013; Ly et al., 2014; Olivares et al., 2017). Furthermore, rodent models inherently possess distinct disparities when juxtaposed with human retinas. Key differences include the lack of a cone-rich macula, different types of photoreceptor cells, and different proportions of these and other cellular subtypes. Given these differences, the unique attributes of human physiology and pathophysiology limit the direct translation of findings derived from animal models to human (Achberger, Haderspeck, et al., 2019).

The advent of stem cell-based models and cell technologies have opened new avenues for basic research and clinical applications. In the retina field, the development of three-dimensional (3D) retinal organoid derived from human induced pluripotent stem cells (hiPSCs) are at the front of these efforts (Osakada et al., 2009; Nakano et al., 2012; Sasai et al., 2012). Retinal organoids largely resemble the human tissue architecture and recapitulate the cellular interactions, which is crucial to the development and function of their *in vivo* counterparts (Meyer et al., 2011; Zhong et al., 2014; Wahlin et al., 2017; Hallam et al., 2018; Achberger, Haderspeck, et al., 2019; Capowski et al., 2019). Organoids are emerging not only to be an effective model in the translational arena but also a valuable tool for understanding human development, physiology, and disease (Kruczek and Swaroop, 2020; McLenachan et al., 2021). In recent years, retinal organoids have been used as models for several purposes such as to identify the mechanisms of inherited retinal degenerative diseases (Zhang et al., 2021), to test advanced therapies based on Adeno-associated virus (AAV) infection for retinal cells (Achberger et al., 2021; Kruczek et al., 2021), for RNA-based therapies (Dulla et al., 2018), to study retinal viral infections such as Severe Acute Respiratory Syndrome Coronavirus 2 (SARS-CoV-2) (Menuchin-Lasowski et al., 2022), and as a source of human cone photoreceptors for cell replacement therapies (Ribeiro et al., 2021). Moreover, retinal organoids are also being explored in novel microphysiological systems (MPS) involving retina-on-chip co-culture approaches (Achberger, Probst, et al., 2019).

To the best of our knowledge, we have established the first 3D human retinal organoid model to study the early molecular mechanisms in DR. We cultured retinal organoids derived from hiPSCs under high glucose conditions to mimic the retinal environment in DR. We observed retinal neurodegeneration, specifically in retinal ganglion and amacrine cells, and an inflammatory response accompanied by the expression of interleukin-1β (*IL-1β*), *VEGF* and secretion of monocyte chemoattractant protein-1 (MCP-1). Additionally, we found increased reactive oxygen species (ROS) levels and activation of the antioxidative stress response and mTOR pathway. Our findings show that our 3D human retinal model successfully replicates several disease features of DRN, serving as an innovative platform for investigating DR mechanisms and pinpointing potential therapeutic targets. This approach offers a compelling preclinical model of early-stage DR, aligning with the 3Rs principle and the call for alternative and non-animal research methodologies. Our methodology is poised to reduce reliance on animal models, enabling data prioritisation for subsequent validation and diminishing risks associated with data translation to human applications.

## Materials and Methods

### Human induced pluripotent stem cell (hiPSC) culture

hiPSC line (IMR90-4, WiCell) derived from a healthy donor and fully characterized was cultured with mTeSR^TM^ Plus Medium (STEMCELL Technologies) on growth factor reduced matrigel-coated 6-well plates (Corning) and used for all experiments in this study. The hiPSCs were routinely passaged using Versene (Gibco, Thermo Fisher Scientific) at a ratio of 1:4-1:6 in mTeSR^TM^ Plus medium supplemented with 10 μM of ROCK inhibitor Y-27632 (Focus Biomolecules) and maintained at 37 °C in a humidified atmosphere containing 5% CO_2_.

### Retinal organoid differentiation of iPSCs

The method for differentiation hiPSCs towards retinal organoids was carried out according to a previously described protocol (Capowski et al., 2019) with slightly modifications to improve the efficiency and the reproducibility in the generation of retinal organoids. Briefly, hiPSCs at 80% of confluence were lifted using Versene (Gibco, Thermo Fisher Scientific) and split at 3000 cells/well in a Nunclon Sphera 96-well U-bottom plate (Thermo Fisher Scientific) using mTeSR^TM^ Plus medium supplemented with 10 μM of ROCK inhibitor (Focus Biomolecules). The cells were grown as aggregates/embryoid bodies for 6 days, doing an adaptation to Neural Induction Medium (NIM) containing DMEM/F12, 1% N2 supplement, 1X Non-Essential Amino Acids, 1X GlutaMax (all from Gibco, Thermo Fisher Scientific) and 2 mg/ml Heparin (Sigma-Aldrich). On day 6, 1.5 nM of Bone Morphogenic Protein-4 (BMP4, Peprotech) was added to fresh NIM, and on day 7 EBs were transferred to growth factor reduced matrigel-coated 6-well plates (Corning). The medium was replaced by half fresh NIM on days 9, 12 and 15. On day 16, the medium was replaced by retinal differentiation medium (RDM) containing DMEM:F12 (3:1), 2% B27 minus vitamin A, 1X Non Essential Amino Acids, 1X GlutaMax, 1X Antibiotic/Antimycotic (all form Gibco, Thermo Fisher Scientific) and changed every 2-3 days. At day 30, optic vesicles were manually dissected using a surgical scalpel (SM65A, Swann-Morton Ltd) under the microscope EVOS XL core (Thermo Fisher Scientific). After dissection, organoids were maintained in suspension flasks (Sarstedt) in 3D-retinal differentiation medium (3D-RDM) containing DMEM:F12 (3:1), 2% B27 minus vitamin A, 1X Non-Essential Amino Acids, 1X GlutaMax, 1X Antibiotic/Antimycotic, 5% FBS, 1:1000 chemically defined lipid supplement (all from Gibco, Thermo Fisher Scientific), 100 μM taurine (Sigma-Aldrich) and 1 μM of all-trans retinoic acid (RA, Sigma-Aldrich) until day 100. After this differentiation stage, organoids were maintained in 3D-RDM and processed for high glucose treatments.

### High glucose treatments

Organoids on day 100 were exposed to different glucose concentrations. Namely, 3D-RDM medium (containing a standard glucose concentration of 19 mM) was supplemented with D-Glucose (Sigma-Aldrich) to a final concentration of 50 mM and 75 mM. D-Mannitol (Sigma-Aldrich) was used as osmotic control (56 mM of Mannitol was added to the 19 mM of glucose already present in the 3D-RDM medium, to obtain an equivalent osmolality as in the highest glucose condition, 75 mM). Control organoids were maintained in regular 3D-RDM medium with 19 mM of glucose. Organoids were maintained for 6 days, with fresh medium exchange every other day. After treatment, organoids were then collected for further analysis.

### Measure of intracellular ROS levels

ROS was measured using the cell permeable fluorogenic probe 2’,7’-Dichlorodihydroflurescein diacetate (DCF-DA, Sigma-Aldrich), widely used to determine the degree of overall oxidative stress. Briefly, after glucose treatment, organoids were incubated in 3D-RDM medium with 20 μM of DFC-DA for 1h at 37°C. Positive control was done using 0.5% H_2_O_2_ for 30 minutes. After incubation, organoids were rinsed once in PBS and plated in chamber slides for live imaging using confocal microscope Zeiss LSM 980 (Zeiss). The mean fluorescence intensity per image was determined using Fiji (Image J) software.

### Glutamine release assay

Glial function was quantified as intracellular conversion of L-Glutamate to L-Glutamine (Mongin et al., 2011). L-Glutamic acid (Sigma) was added at 3 mM in GlutaMax-free 3D-RDM medium at the third day of glucose treatment. Medium culture samples were collected at day 6, centrifugated at 200 *g*, 4 °C for 10 min and supernatants were collected for glutamine analysis. Glutamine quantifications were measured using Cedex Bio analyser 7100 (Roche), and concentrations were normalized by the protein content of each sample (mmol/L/µg).

### Histology and Immunofluorescence

Retinal organoids were fixed in 4% Paraformaldehyde phosphate-buffer solution (PFA, Sigma) for 20 minutes at room temperature. Washed twice with phosphate-buffered saline (PBS) and incubated in 10% for 1h, 20% for 1h, and 30% sucrose solution overnight at 4 °C. Retinal organoids were placed into cryogenic moulds and immersed in optimum cutting temperature (OCT Cryomatrix, Fisher Scientific). About 12 μM slices were sectioned on a Leica CM3050 S cryostat (Leica). Cut sections were permeabilized with 0.05% Triton-X-100 (Sigma-Aldrich) in PBS for 15 minutes and were blocked in 10% Donkey Serum (DS) in PBS for 1 hour. Primary antibodies were incubated in blocking solution at 4 °C overnight. The following primary antibodies were used: mouse anti-AP2α (1:100, sc-12726, Santa Cruz Biotechnology), goat anti-BRN3a (1:100, sc-31989, Santa Cruz Biotechnology), goat anti-OTX2 (1:100, AF1979, R&D Systems), rabbit anti-Vimentin (1:200, 5741, Cell Signalling), Alexa Fluor 568 Phalloidin (1:400, A12380, Invitrogen). Sections were then washed three times with PBS followed by incubation of secondary antibodies for 1 hour at room temperature. The following secondary antibodies were used (all from Invitrogen and used at 1:1000 dilution): Donkey anti-goat 488 (A11055), Donkey anti-mouse 488 (A21202), Donkey anti-rabbit 488 (A21206). Sections were then incubated with 4’, 6-diamidino-2-phenylindole (DAPI, 1 μg/ml, Sigma-Aldrich), washed twice with PBS and mounted with Vectashield mounting medium (Vector Laboratories). Confocal images were acquired using Zeiss LSM 980 (Zeiss) and images were processed using Image J Software. For the quantitative analysis of the number of retinal populations (specifically RGCs, amacrine cells and photoreceptor progenitors) positive cells were expressed relative to the total number of DAPI stained nuclei (at least five sections per condition were analysed from three independent experiments). For fluorescence analysis and quantification, Vimentin intensity was measured using Image J Software (at least five sections per condition were analysed from three independent experiments).

### Fluoro-Jade C staining

Fluoro-Jade C is an anionic fluorochrome commonly used for identifying degenerating neurons in the brain and in the retina regardless the neurotoxic insult or the mechanism of cell death (Schmued et al., 1997; Chidlow et al., 2009; Oshitari et al., 2011). Fluoro-Jade C labelling was performed on retinal organoids cryosections using the Fluoro-Jade® C staining kit (Biosensis) and following the manufacturer’s instructions. After co-incubation of Fluoro-Jade C and 4,6-diamidino-2-phenylindole (DAPI), slides were air dried for 10 minutes, cleared in xylene and mounted with Entellan^TM^ (Sigma-Aldrich) mounting media. Samples were analysed in the confocal microscope Zeiss LSM980 (Zeiss) and Fluoro-Jade C^+^ cells of each section were evaluated. For the quantitative analysis, the number of Fluoro-Jade C^+^ cells were expressed relative to the total number of DAPI stained nuclei (at least five sections per condition were analysed from three independent experiments).

### Western blot analysis

Pools of 2 to 4 organoids were resuspended in cold lysis buffer (Cell Signalling Technology) supplemented with protease and phosphatase inhibitor cocktails (Roche) and sonicated twice for 3 seconds at 10% intensity in the Branson digital Sonifier SFX 150 (Emerson). Lysates were then centrifuged at 7500 *g* for 10 minutes at 4 °C, and supernatant were collected for protein quantification using the Pierce BCA protein assay kit (Thermo Scientific) following the manufacturer’s instructions. A total of 20 μg of protein from each lysate were resolved on 10% or 12% sodium dodecyl sulphate – polyacrylamide gels (SDS-PAGE) and subsequently transferred to nitrocellulose membranes (Bio-Rad Laboratories). Membranes were blocked using 5% non-fat dry milk or 5% bovine serum albumin (BSA) (Sigma-Aldrich) in Tris-buffered saline (TBS) (50 mM Tris, 150 mM NaCl, pH = 7.6) containing 0.1% Tween-20 (Sigma-Aldrich) (TBS-T). Primary Antibodies were incubated in blocking solution overnight at 4°C. The following primary antibodies were used: rabbit anti-Glutamine Synthetase (1:1000, NB110-41404, Novus Biologicals), mouse anti-Catalase (1:500, sc-271803, Santa Cruz Biotechnology), mouse anti-SOD1 (1:1000, sc-31989, Santa Cruz Biotechnology), rabbit anti-SOD2 (1:1000, ab13533, Abcam), rabbit anti-phospho-Akt Ser473 (1:1000, 9271, Cell Signaling Technology), rabbit anti-Akt (1:1000, 9272, Cell Signaling Technology), rabbit anti-phospho-S6 Ribosomal protein Ser235/236 (1:1000, 4858, Cell Signaling Technology), rabbit anti-S6 Ribosomal Protein (1:1000, 2217, Cell Signaling Technology), mouse anti-β-Actin-peroxidase (1:25000). After washing with TBS-T, the appropriate HRP-conjugated secondary antibody was added (1:5000 in blocking buffer) for 2 h at room temperature. Secondary antibodies used: donkey anti-rabbit HRP (NA934, Cytiva) and sheep anti-mouse HRP (NA931, Cytiva). Antibody binding was detected using chemiluminescence ECL Prime Western Blotting Substrate (Cytiva) and images were acquired on ChemiDoc Touch (Bio-Rad Laboratories). The acquired images were processed and quantified using Image Lab software (Bio-Rad laboratories) and the protein of interest were normalized using β-Actin as a loading control.

### RNA extraction and Quantitative real-time PCR

Total mRNA from 2 to 4 organoids were isolated using the RNeasy mini kit (Qiagen) according to manufacturer’s protocol. 1 μg of mRNA was reverse transcribed into cDNA using Superscript II reverse transcriptase kit (Invitrogen). The cDNA samples were used for quantitative PCR in the LightCycler 96 system (Roche) using the FastStar Essential DNA Green Master (Roche) following the manufacturer’s instructions. Primers pairs were designed using Primer-BLAST (NCBI) and synthesized by Thermo Fisher Scientific. The following primer sequences were used (forward/reverse): *AKT*, (5’-TGATCACCATCACACCACCT-3’ / 5’-CTGGCCGAGTAGGAGAACTG-3’); *GPX, (5’-*TGGGCATCAGGAGAACGCCA-3’/ 5’-GCGTAGGGGCACACCGTCAG-3’); *HIF1⍺*, (5’-CAGTCGACACAGCCTGGATA-3’/5’-GCGGCCTAAAAGTTCTTCTG-3’); *IL1B,* (5’-GTTTCTCTGCAGAAAGAGGC-3’/5’-AATGCCAGAGATGCATTGG-3’); *mTOR*, (5’-CTGGTTTCACCAAACCGTCT-3’/5’-GCACGACGTCTTCCAGTACC-3’); *SOD1,* (5’-TGGCCGATGTGTCTATTGAA-3’/5’-ACCTTTGCCCAAGTCATCTG-3’); *SOD2*, (5’-TGGTTTCAATAAGGAACGGG-3’/5’-GAATAAGGCCTGTTGTTCCT-3’); *VEGF*, (5’-CCTTGCTGCTCTACCTCCAC-3’/5’-ATGATTCTGCCCTCCTCCTT-3’; *β-Actin*, (5’-GAAGATCAAGATCATTGCTCCTC-3’/5’-ATCCACATCTGCTGGAAGG-3’). The expression levels were normalized to the housekeeping β-Actin and fold change was calculated using the 2^-ΔΔCt^ method.

### Enzyme-Linked Immunosorbent Assay (ELISA)

MCP-1 levels secreted by organoids were assessed using the Human MCP-1 Standard TMB ELISA development kit (Peprotech) according to the manufacturer’s protocol. Briefly, cell supernatants were plated in duplicates and incubated with MCP-1 detection antibody and HRP-streptavidin conjugate. After stopping colour development reaction, the absorbance was measured at 450 nm using a Synergy HT microplate reader (Agilent) and normalized by the protein content of each sample ((ng/ml)/μg of protein).

### Data analysis and statistics

Statistical analyses between treatment groups and controls were performed using Prism 8 (GraphPad Software). Data are presented as the mean±SD and statistical tests were conducted using the One-way ANOVA followed up by Tukey’s multiple comparisons tests. All statistical comparisons were performed on data from ⩾ 3 biologically independent experiments. Significance is shown as \**p*<0.05, \*\**p*<0.01, \*\*\**p*<0.001.

## Results

### High glucose treatment induces neurodegeneration in retinal organoids

For this study, retinal organoids were generated from hiPSC line IMR90-4 according to the three-stage differentiation protocol described by Capowski et al. (Capowski et al., 2019). We used retinal organoids differentiated for 100 days. At this stage, these organoids contain pivotal retinal cells affected in DR. Specifically: *i*) most cells of the neuroretina are present, such as RGCs located in the basal region; *ii*) Photoreceptor progenitors are found in the apical layers, and starburst amacrine cells are also identified (Capowski et al., 2019; O’Hara-Wright and Gonzalez-Cordero, 2020); *iii*) Müller glia progenitors span the entire width of the organoid, mirroring the distribution of Müller cells in the human retina (Eastlake et al., 2019); *iv*) the 100-day organoids display a complete lamination of the inner retina (Capowski et al., 2019). Given these features, retinal organoids at 100 days of differentiation offer a physiologically relevant model for disease modeling (Mahato et al., 2022), drug testing, and even cell transplantation (Eastlake et al., 2019). First, we exposed retinal organoids with 100 days of differentiation to either 19 mM of glucose (standard glucose concentration in the 3D-RDM medium) or high glucose concentrations (50 mM and 75 mM) for 6 days. The nuclei staining with DAPI of cross-sections revealed a significant increase in the percentage of pyknotic nuclei on the organoids exposed to 75 mM of glucose in comparison with control and 50 mM of glucose (Fig. 1A, images a-c and 1B). Furthermore, Mannitol treated organoids did not show significant differences in the number of pyknotic nuclei in comparison with the control condition (Fig.1A, image d and 1B). In parallel, we evaluated neurodegeneration induced by high glucose conditions by assessing the percentage of positive cells for Fluoro-Jade-C (FJ-C) staining, which is considered a reliable marker of degenerating immature and mature neurons, including apoptotic, necrotic, and autophagic cells (Ikenari et al., 2020). We observed a significantly increased percentage of FJ-C positive cells in cross-sections of organoids exposed to 75 mM compared to the control condition (Fig. 1A, images e-h and 1C).

**Figure 1:**
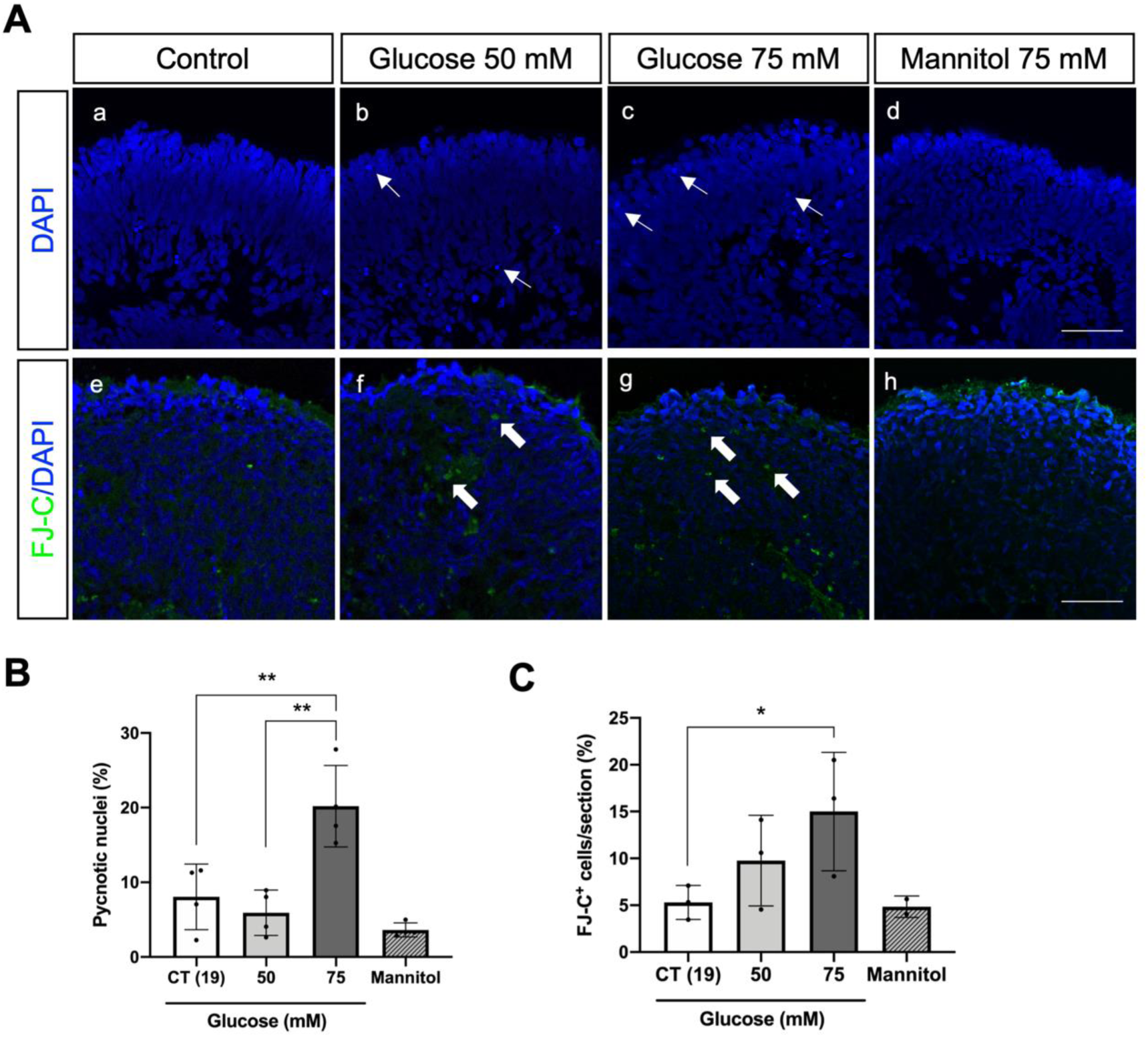
High glucose treatment induces neurodegeneration in day 100 retinal organoids. **A**) Cross-sectional images of retinal organoids with DAPI nuclei staining (in blue) showing pyknotic nuclei (a-d) and (e-h) represent cross-sectional images of retinal organoids with Fluoro-Jade-C (FJ-C) staining (in green). White arrows indicate pyknotic nuclei (thin) and degenerating neurons (thick). Scale bar: 50 µm. **B**) Quantification of pyknotic nuclei after high glucose treatment. **C**) Quantification of FJ-C positive cells (FJ-C^+^) after high glucose treatment. Statistical results are the mean values±SD of 2-3 organoids from at least 3 independent differentiations (\**p* value<0.05 and \*\**p* value<0.01).

Overall, these data show that the exposure of the retinal organoids to high glucose induces cell death and neurodegeneration, as reported in the early stages of DR in both animal models and in the eyes of diabetic patients (Barber et al., 1998; Sohn et al., 2016; Chakravarthy and Devanathan, 2018; Simó et al., 2018; Zafar et al., 2019; Channa et al., 2021; Soni et al., 2021).

### High glucose treatment induces retinal ganglion and amacrine cell loss in retinal organoids

Several studies performed in human retinal sections from diabetic patients have shown that RGCs are amongst the most affected retinal cells in DRN. Amacrine and photoreceptor cell death has also been observed in retinal sections (Barber et al., 1998; Carrasco et al., 2007; Kern and Barber, 2008; Garcia-Ramírez et al., 2009). We evaluated how high glucose exposure affected the different retinal cell types of the retinal organoids. AP2α belongs to a family of AP2 transcription factors and is commonly used as a marker of amacrine cells. However, AP2α is also known to be expressed in developing horizontal cells (Bassett et al., 2012; Hicks et al., 2018), as is the case of retinal organoids at day 100 of differentiation. Indeed, the number of AP2α positive cells per section decreased by 50% in retinal organoids exposed to high glucose conditions, indicating a reduction of amacrine and horizontal cells (Fig. 2A, images a-c and 2B). Moreover, the number of BRN3a positive cells (RGCs) per section presented a significant 2-fold decrease at higher glucose conditions (75mM) (Fig. 2A, images d-f and 2C), while the number of OTX2 positive cells (photoreceptor progenitor cells) has not changed (Fig. 2A, images g-i and 2D). This data reinforces that treating retinal organoids for 6 days with high glucose concentrations is enough to reproduce the loss of specific populations observed in DRN.

**Figure 2:**
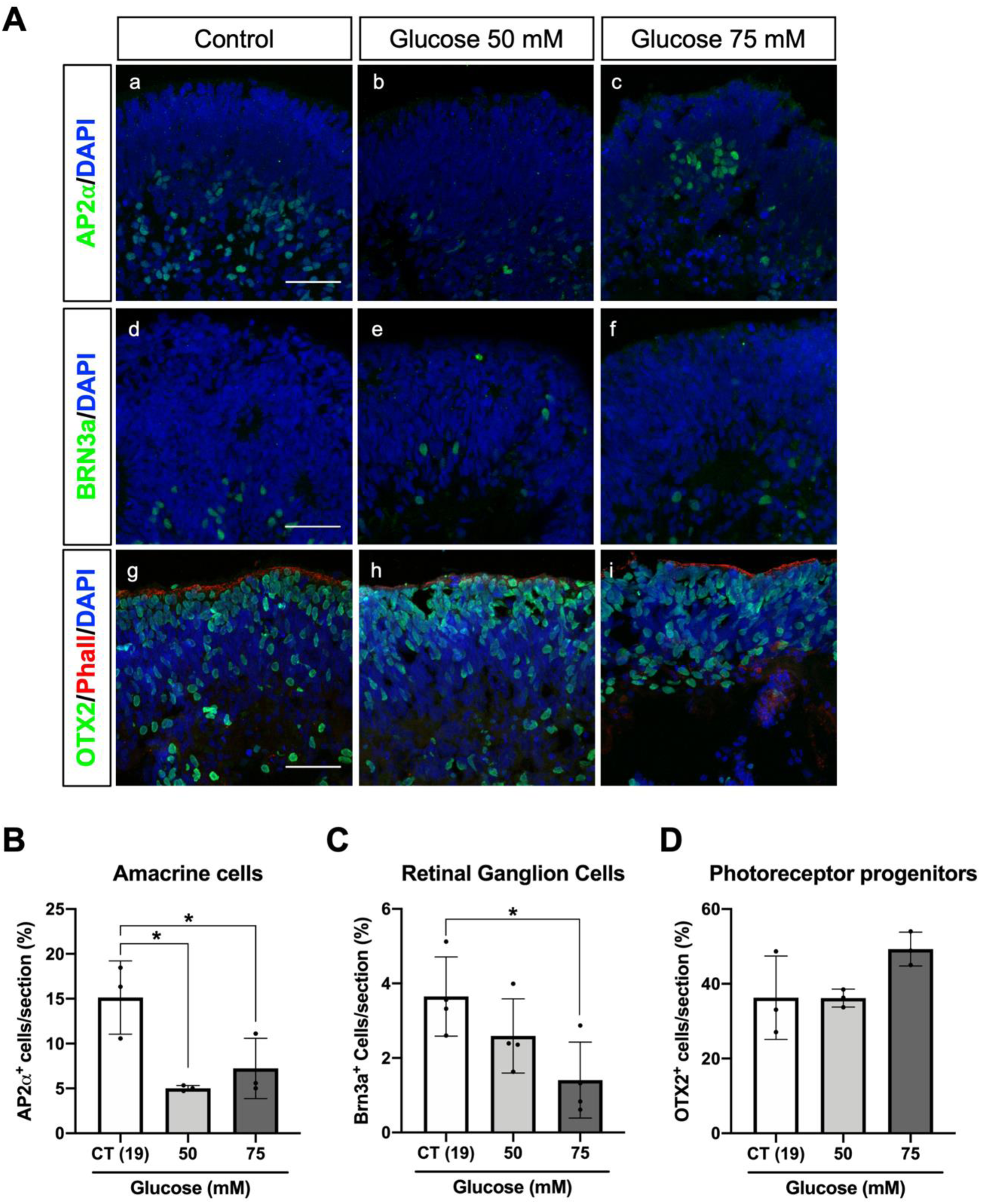
High glucose treatment induces amacrine and retinal ganglion cell loss. **A**) Immunofluorescence analysis of organoids subjected to high glucose (a-c) staining of AP2⍺ (amacrine cells), (d-f) staining of BRN3a (retinal ganglion cells) and (g-i) staining of OTX2 (photoreceptor progenitors) and Phalloidin (outer limiting membrane). Organoids sections were counterstained with DAPI. Scale bar: 50 µm. **B)** Quantification of AP2⍺-positive cells counted per field of view. **C)** Quantification of Brn3a-positive cells counted per field of view. **D)** Quantification of OTX2-positive cells counted per field of view. Data represent mean±SD of at least 3 independent differentiations (\**p* value<0.05).

### High glucose treatment induces inflammation in retinal organoids

Müller cells, the principal glial cells in the retina, are thought to be a major source of inflammatory factors in DR (Mizutani et al., 1998; Rübsam et al., 2018). Müller cells express and secrete several growth factors and cytokines that alter the function and survival of retinal neurons and capillary cells. These cells could be responsible for production and secretion of several inflammatory mediators including VEGF, MCP-1, tumor necrosis factor-α (TNF-α), IL-1β, interleukin-6 (IL-6) (Rübsam et al., 2018). Functional Müller cells contribute to the removal of the extracellular glutamate and production of glutamine essential for retinal neurons. Under pathological conditions, such as glaucoma or ischemia, the dysregulation of Müller cells showed a decrease in the glutamate uptake and the glutamine release caused by the impairment of glutamine synthetase (Nishiyama et al., 2000; Kruchkova et al., 2001; Moreno et al., 2005). In contrast, in DR and optic nerve crush, no changes or a slight increase of glutamine synthetase is observed (Mizutani et al., 1998; Lo et al., 2001). Our results showed, that at day 100, retinal organoids present Müller glia progenitors as indicated by immunostaining of cross-sections with the corresponding marker vimentin (Fig. 3A). Interestingly, vimentin immunostaining signal increases significantly and cytoskeleton morphology changes are observed under high glucose treatment (Fig. 3A and B). Additionally, the protein levels of glutamine synthetase were 4-fold increase in organoids exposed to 50 mM compared to the control conditions (Fig. 3C) suggesting an increase in glial function. Moreover, the levels of glutamine release were significantly increased in organoids exposed to 75 mM of glucose compared to the control conditions (from approximately 0.15 to 0.25 mmol/L/ug) (Fig. 3D). The increase of glutamine synthetase may be attributed to an increase of the defence against oxidative stress and as a proactive response to safeguard against neuronal degeneration under high glucose conditions (Gorovits et al., 1997). Furthermore, considering the pivotal role of Hypoxia-Inducible Factor 1*α* (*HIF-1α*) in regulating cellular oxygen homeostasis and aiding adaptation to hypoxia during DR pathogenesis - along with its role in the expression of pro-inflammatory cytokines and VEGF (Zhong et al., 2012) - we decided to assess the expression of *HIF-1α* and *VEGF*. Our results showed that *HIF-1α* expression does not change. However, *VEGF* expression was almost a 2-fold increase in retinal organoids under high glucose treatment (Fig. 3E). In addition, the pro-inflammatory cytokine *IL-1β* expression was increased when treated with 75mM of glucose. (Fig. 3F). Moreover, MCP-1 secretion is known to be produced by glial cells in diabetic patients (Taghavi et al., 2019). Interestingly, we observed increased secretion of MCP-1 at 50 mM of glucose (Fig. 3G). Altogether, our results show that under our experimental design, retinal organoids at day 100 already present markers of gliosis and inflammation, which are essential features of DRN (Tang and Kern, 2011; Rübsam et al., 2018).

**Figure 3:**
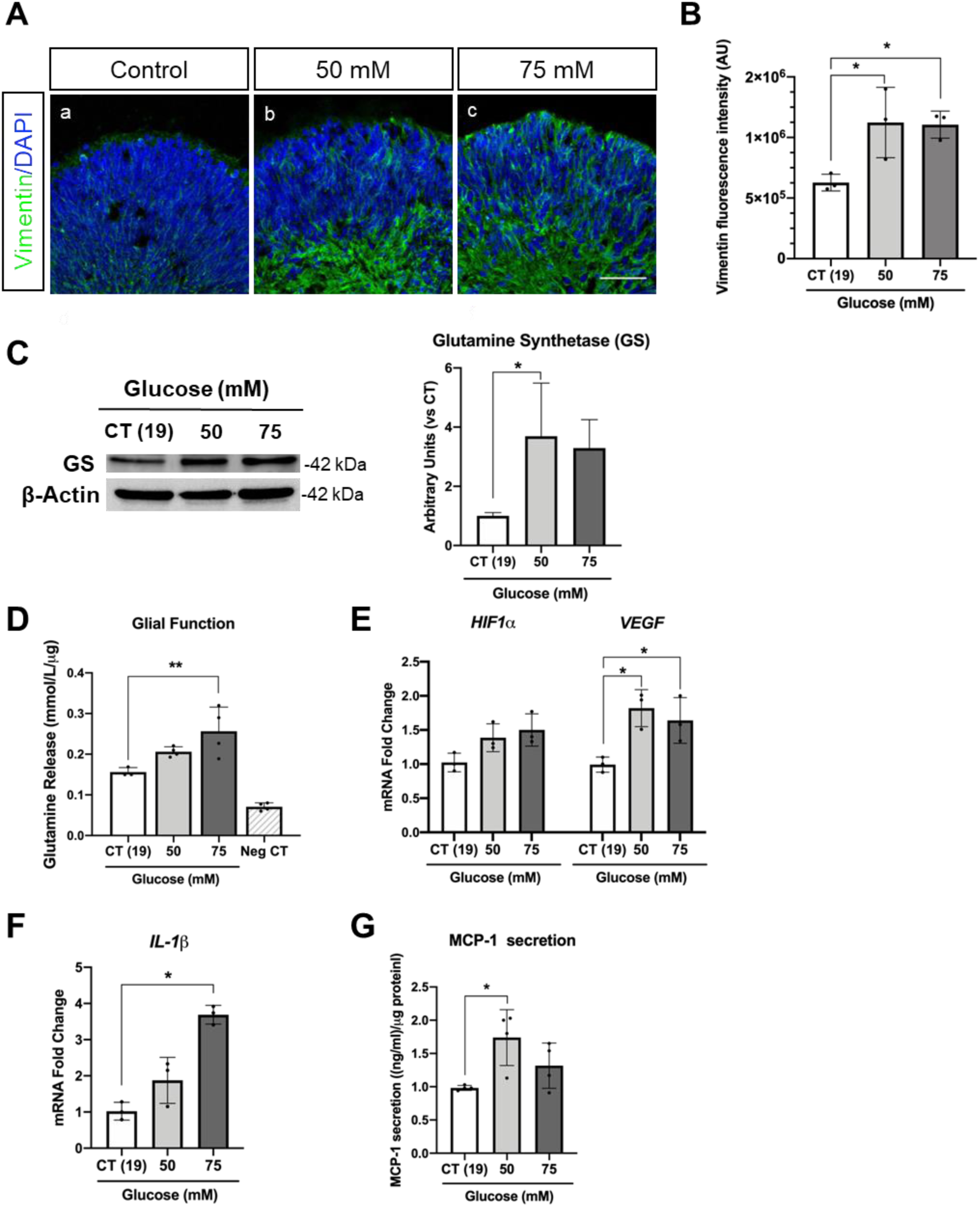
High glucose treatment induces an inflammatory response in retinal organoids. **A**) Immunofluorescence staining of Vimentin (a-c) counterstained with DAPI (blue). Scale bar: 50 µm. **B)** Confocal image quantification of Vimentin fluorescence intensity. **C**) Western blot analysis of Glutamine Synthase (GS) (left panel) and correspondent densitometry analysis of GS normalized by β-Actin (right panel). **D)** Glutamine release in the culture medium after high glucose treatment, reflecting the conversion of glutamate to glutamine by the glial cells. **E**) Expression of *HIF1⍺* and *VEGF* mRNA. **F**) Expression of the pro-inflammatory cytokine *IL-1β* mRNA. **G)** MCP-1 secretion levels in the culture medium measured by ELISA. Results represent the mean±SD of at least 3 independent differentiations (\**p* value<0.05 and \*\**p* value<0.01).

### High glucose treatment induces oxidative stress in retinal organoids

Oxidative stress stands as a defining feature of DR. Under the chronic influence of elevated glucose levels, there is an increased production of ROS spurred by the activation of the secondary pathways such as the polyol and hexosamine pathways. Furthermore, the overproduction of AGEs concurrently increases ROS production, thereby establishing a self-perpetuating cycle (Hammes, 2018; Cecilia et al., 2019). To determine whether oxidative stress was induced in the experimental conditions used in this study, we first evaluated the production of ROS levels assessed by the 2’-7’-dichlorofluorescin diacetate (DCF-DA) probe. Using the fluorescence intensity of DCF-DA as captured by confocal imaging, we observed a notable increase in intracellular ROS production in organoids exposed to high glucose conditions (75 mM) when compared to both the control and the 50 mM conditions. (Fig. 4A). Additionally, we examined the gene expression and protein levels of pivotal enzymes central to the primary defense against oxidative stress, including superoxide dismutases (SODs), Catalase, and Glutathione Peroxidase 1 (GPX1). Notably, while our results revealed no changes in *SODs* expression levels (Fig. 4B), there was a significant increase in the protein levels of SOD1 and SOD2 in retinal organoids under high glucose conditions. This suggests that post-transcriptional regulation of these enzymes is taking place (Fig. 4C-E). Moreover, no changes were observed in the expression of *GPX1* under high glucose conditions (Fig. 4B). However, there was a notable increase in Catalase protein levels at 50 mM conditions (Fig. 4C, F). Collectively, these findings suggest that an antioxidant response is activated in retinal organoids exposed to high glucose concentrations, likely to counteract the effects of ROS.

**Figure 4:**
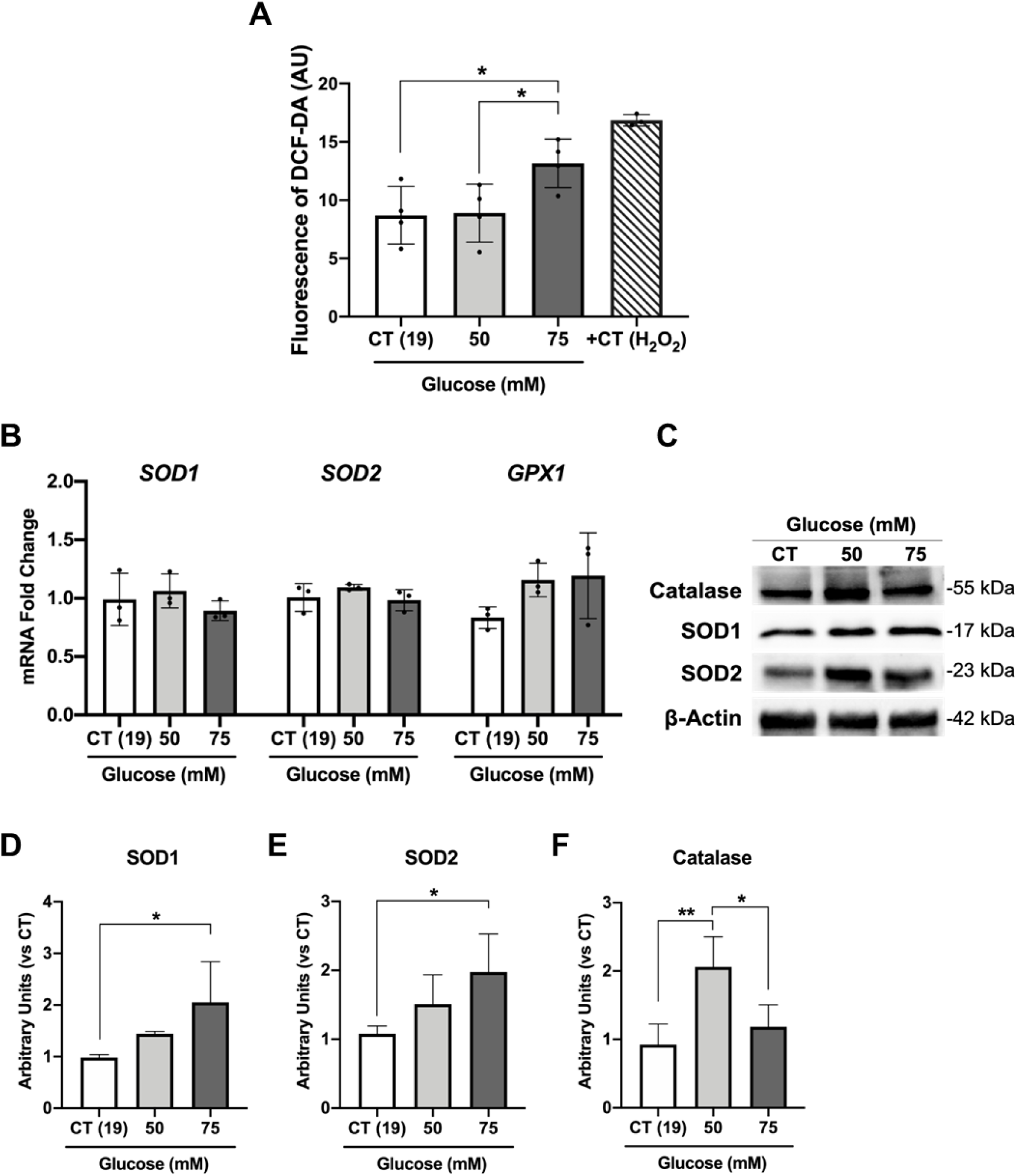
High glucose treatment affects the antioxidant response in organoids. **A)** Fluorescence intensity of 2,7-dichlorofluorescein diacetate (DCF-DA) probe for detection of reactive oxygen species in organoids after high glucose exposure (n=4). **B)** Expression of *SOD1*, *SOD2* and *GPX1* mRNA levels. **C**) Western blot analysis of catalase and superoxide dismutase 1 and 2 (SOD1 and SOD2). **D-F**) Densitometry analysis of SOD1, SOD2 and catalase normalized by β-Actin. Data represent the mean±SD of at least 3 independent differentiations (\**p* value<0.05 and \*\**p* value<0.01).

### High glucose treatment induces mTOR pathway in retinal organoids

The phosphatidylinositol 3-kinase (PI3K)/AKT/mammalian target rapamycin (mTOR) signaling pathway is a well-known central player on large number of biological events related to cell growth, division and metabolism (Szwed et al., 2021). Moreover, dysregulation in mTOR is associated with various diseases such as obesity, diabetes, cancer and neurological diseases (Saxton and Sabatini, 2017). In DR, there is significant evidence that the PI3K/AKT/mTOR signaling pathway is connected to various mechanisms associated with disease progression, including oxidative stress, inflammation, hypoxia, angiogenesis and proliferation (Safi et al., 2014; Ahmad, 2015). Therefore, we evaluated the expression levels of *mTOR* and *AKT.* A marked elevation in the expression of both genes was observed in retinal organoids subjected to 75 mM glucose, in contrast to the control (19 mM) and 50 mM treatment (Fig. 5A). Neither the AKT protein nor its S473 phosphorylation levels displayed significant changes under the conditions tested (Fig. 5B, C). Nevertheless, the phosphorylation levels of the mTOR’s primary downstream effector, the ribosomal S6 kinase (S6) at Ser235/236, showcased a significant increase in retinal organoids at 75 mM glucose when compared to the control and 50 mM treatment (Fig. 5B, D).

**Figure 5:**
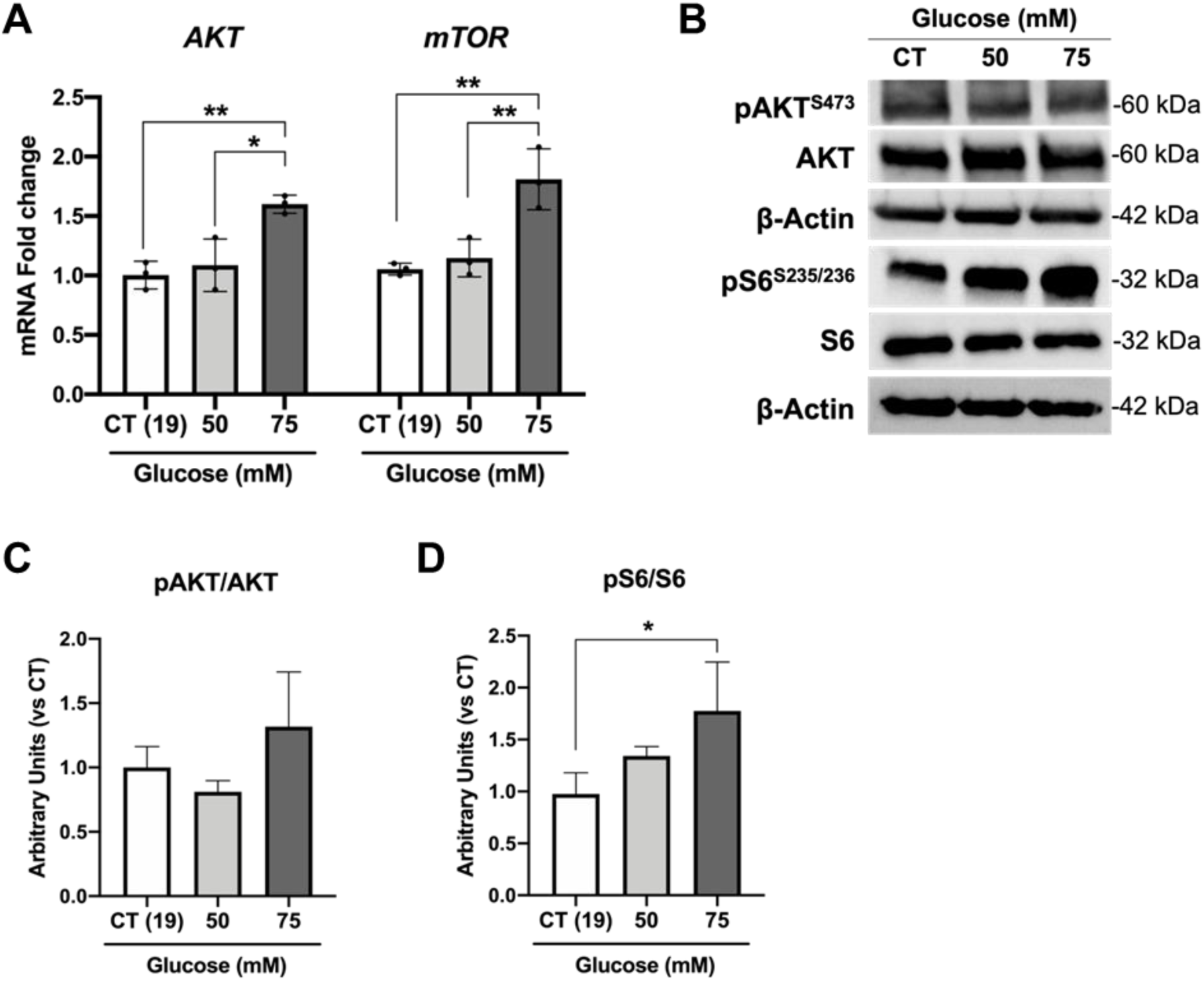
High glucose treatment effects on mTOR signalling pathway. **A)** Expression levels of *AKT* and *mTOR* mRNA levels after high glucose treatment. **B)** Western blot analysis of phospho-AKT^S473^, AKT, phospho-S6S^235/236^ and S6 protein levels normalized by β-Actin. **C**) Densitometry of pAKT/AKTlevels after glucose treatment. **D)** Densitometry of pS6/S6 protein levels after high glucose conditions. Data represent mean±SD of at least 3 independent differentiations (\**p* value<0.05 and \*\**p* value<0.01).

Together, these findings highlighted the activation of mTOR signaling pathway in retinal organoids exposed to high levels of glucose.

## Discussion

The study and comprehension of any human disease are intimately dependent on the available experimental models. Both *in vitro* and *in vivo* DR models have been instrumental in clarifying the molecular and cellular mechanism involved but also to identify new therapies. Nevertheless, every existing DR model possesses its own set of advantages and drawbacks. Retinal organoids are emerging as powerful models to study retina development, diseases, toxicology, and new therapies (Kruczek and Swaroop, 2020; Zhang et al., 2021). They can be derived from human adult or embryonic stem cells (ESCs) or hiPSCs obtained from healthy individuals or patients exhibiting specific diseases. In advanced stages of differentiation, these 3D structures effectively mimic all neuroretina cell types, including Müller cells and astrocytes, presenting them in an organized stratified layout reminiscent of the human retina. Moreover, these organoids can develop mature photoreceptors that are not only photosensitive but also capable of phototransduction, allowing advanced electrophysiological research (Artero Castro et al., 2019; Cowan et al., 2020). A shared trait among various organoid types, retinal ones included, is the absence of vascularization and microglia. Additionally, due to limited nutrient availability, there’s an observed attrition of cells in the innermost retinal layer of the organoids. This phenomenon becomes particularly pronounced in the RGCs during extended differentiation periods, specifically beyond 120 days (Achberger, Haderspeck, et al., 2019).

In the present study, we established a retinal organoid model for DR, with a particular focus on DRN. Clinical studies based on optical coherence tomography in diabetic patients have shown that these patients often display a progressive loss of the ganglion cell and nerve fiber layers (Sohn et al., 2016). Additionally, these patients can experience vision changes, even in the absence or presence of minimal DR (Zafar et al., 2019). Also, studies in human retinas further suggests that changes in neuronal cells, such as cell death and axon degeneration, occur before the vascular abnormalities arise (Sohn et al., 2016; Channa et al., 2021). These findings are supported by both *in vitro* and *in vivo* studies (Oshitari et al., 2011). Given these findings, we believe that retinal organoids present represent a valuable model for exploring the initial stages of DR, dissecting DRN without the interference of the vascular system. Moreover, we limited our study to 100-day-old organoids to circumvent the potential loss of RGCs, which is typical of advanced differentiation stages.

Our *in vitro* experimental design replicates the molecular features observed in DR patients by adjusting the glucose concentrations in retinal organoids medium to concentration between 50 and 75 mM. While the glucose concentrations used in our study do not directly correspond to the serum levels found in either healthy or diabetic patients, it is important to note that our standard culture medium has a basal glucose concentration of 19 mM (control conditions). This reflects the high metabolic rate of neuronal cells in comparison to other cell types. Notably, the neuroretina is the neuronal tissue with the highest energy demand (Country, 2017). Photoreceptors, which predominantly absorb glucose in the retina, obtain their glucose supply from the choroidal blood (Kanow et al., 2017). This glucose is channeled through the basolateral and apical membranes of the RPE before reaching the photoreceptors (Kanow et al., 2017; Li et al., 2020). Interestingly, under the conditions we tested, photoreceptor progenitors did not show significant loss due to the high glucose concentrations. This might suggest that either a more extended treatment is necessary to observe alterations, or their metabolic requirement on glucose enables them to cope with these conditions. The loss of photoreceptors in DR might be an indirect result of other cells loss or metabolic dysregulation of the RPE supporting cells.

Supplementing media with glucose is a classical approach used across various *in vitro* models. For example, dorsal root ganglion neurons required 25 mM D-glucose in medium for optimal survival (Peeraer et al., 2011). In other studies, concentrations as high as 55 mM were used to evaluate the effects of high glucose (Russell et al., 1999, 2002; Vincent et al., 2005; Fu et al., 2012). Several other models also used glucose concentrations ranging between 30 to 55 mM, or even higher, such as 3D *in vitro* model of the human cornea (Deardorff et al., 2018), Zebrafish embryos (Lee and Yang, 2021), rat retinal explants (Oshitari et al., 2014; Wang et al., 2017), primary cultures of rat hippocampal neurons (Gaspar et al., 2010), primary cultures of rat retinal neural cells (Baptista et al., 2015), rat retinal endothelial cells (Castilho et al., 2012). Also, the duration of these treatments varies widely across studies, lasting from 2 hours to 42 days (in the case of 3D *in vitro* model of the human cornea). This reflects different study aims, from short-term or long-term high-glucose exposure, acute versus chronic conditions.

In diabetic patients, plasma glucose concentrations can soar to levels between 200 to 300 mg/dL (11.1 mM to 16.6 mM). In contrast, healthy individuals typically have fasting blood glucose levels of 99 mg/dL or lower (< 5.5 mM) (accordingly with the Centers for Disease Control and Prevention, CDC). These changes in concentration are similar in proportion to the ones we used in our study, as the high glucose concentrations we tested were 2.5 to 4 times higher than in control.

Through glucose media supplementation, we have demonstrated that retinal organoids exposed to high glucose conditions can recapitulate several characteristics observed in the early stages of DR, particularly in DRN. In these high glucose conditions, we noted not only cell degeneration and an increase in pyknotic nuclei but also a decrease in the number of amacrine and retinal ganglion cells.

Several studies have pinpointed RGCs are the main population affected in the early stages of DR (Kern and Barber, 2008; van Dijk et al., 2012). The death of RGC, axonal degeneration, and ultimately optic nerve degeneration, is central in the retinal neuropathy observed in early DR. While several factors are believed to contribute to RGC malfunction and subsequent death, inflammation and oxidative stress are considered paramount (Potilinski et al., 2020). In our model, we observed clear indications of glial reactivity and inflammation. Under retinal stress, both glial fibrillary acid protein (GFAP) and Vimentin (ubiquitously expressed in retinal glial cells) are well-known sensitive markers for retinal gliosis (Okada et al., 1990; Lewis and Fisher, 2003). Since organoids differentiated at day 100 do not express GFAP (Capowski et al., 2019), mainly due to the absence of mature astrocytes, we detected alterations on the morphology and immunoreactivity of Vimentin, known to be expressed in Müller glia progenitors (Eastlake et al., 2019) in response to high glucose treatments. Noticeably, 6 days of high glucose treatment were also sufficient to induce an increase of MCP-1, *IL-1β* and *VEGF* by glial progenitor cells. According to previous studies, in the early phases of DR, *VEGF* expression and release is increased, which is believed to be the retina’s protective response to prevent cell damage (Amin et al., 1997; Mathews et al., 1997). At this stage, VEGF is a prosurvival agent rather than a proangiogenic factor. However, its role evolves as the disease progresses and VEGF levels remain high. VEGF production by Müller cells is essential for the inflammation and vascular leakage observed in DR, as evidenced in a Müller cell VEGF knockout mouse model (Wang et al., 2010).

Increased production of ROS is a well-studied consequence of high glucose levels, and it is known to be promoted by several metabolic pathways (polyol, hexosamine, AGE and PKC pathways) (Hammes, 2018; Wu et al., 2018; Cecilia et al., 2019). In our experimental setup we also observe increased ROS levels together with an increase in the protein levels of Catalase, SOD1 and SOD2, indicating that antioxidant response is being activated. However, this increased protein levels cannot be translated into an effective ROS prevention. Indeed, it is speculated that post-translational modifications such as O-glycosyl-N-acetylation (O-GlcNAc) and glycation, that are potentiated in hyperglycaemia and diabetes, can affect antioxidant enzymes activity which was found to be decreased in DR animal models and in patients (Cecilia et al., 2019; Bokhary et al., 2021). Interestingly, in a Müller cell line, ROS production in response to high glucose was observed to be dependent on mTOR activity (Kida et al., 2021). The mTOR pathway oversees various biological processes, including protein synthesis, cell proliferation, autophagy, metabolism, and cell survival. In our experiments, exposure to high glucose resulted in increased mTOR activity, correlating with a rise in pS6 levels, a key effector in the mTOR pathway. Studies *in vitro* and *in vivo* showed that mTOR signaling is amplified in streptozotocin (STZ)-injected hyperglycaemic rats and in cultured Müller cells under high glucose conditions (Wei et al., 2016; Kida et al., 2021). While our findings suggest a significant role for mTOR in DR, comprehensive studies are required to fully understand its importance within the disease’s context. Future research should assess whether the molecular and cellular changes observed after 6 days of high glucose exposure can be reversed. It would also be insightful to study the effects of prolonged high glucose treatments, either at the same or reduced concentrations, on the neuroretina.

## Conclusions

We believe that creating a DRN model using human cells can provide a valuable platform for drug screening, in addition to rodent models. This approach could be more relevant for translational research. Additionally, both virus-based and non-virus-based gene therapy methods can be investigated using these 3D models, as evidenced by the successful infection of retinal organoids with AAVs in recent studies (Quinn et al., 2019; Garita-Hernandez et al., 2020; Achberger et al., 2021).

Our 3D retinal model for DR opens new possibilities to study the mechanisms behind this disease. Organoids are a valuable tool to mimic disease, showing several features of DR that are independent of vascularization problems. Long periods of high glucose and more mature organoids that include other retinal cell types, are experimental strategies that can be further explored. Also, establishing more complex systems with vascularization and microfluidics to explore into the vascular mechanisms involved in disease progression presents an interesting approach.

We are confident in the potential of this models because *i*) it uses human cells, yielding reliable results and promoting translational research; *ii*) it offers accessibility, as a significant quantity of organoids can be produced compared to the limited number of eyes available from murine models; *iii*) it can reduce the reliance on animal testing.

## Author Contributions

L. de Lemos developed the model, performed the characterization of the model and the data analysis. P. Antas, I.S. Ferreira, I.P. Santos, C.M. Gomes and C. Brito contributed to the model characterization. L. de Lemos and S. Tenreiro wrote the article. P. Antas, C.M. Gomes, C. Brito and M.C. Seabra revised the article. S. Tenreiro got funding and supervised the project.

## Data availability statement

The data that support the findings of this study are available from the corresponding author upon reasonable request.

## Acknowledgments

This research was funded by Fundação para a Ciência e Tecnologia (FCT) / Ministério da Ciência, Tecnologia e Ensino Superior project PTDC/MED-PAT/29656/2017, iNOVA4Health (UIDB/04462/2020 and UIDP/04462/2020), and by the Associated Laboratory LS4FUTURE (LA/P/0087/2020). CMG was recipient of individual PhD fellowship funded by FCT (UUI/BD/151253/2021).

## Correspondence address

Sandra Tenreiro, iNOVA4Health, NOVA Medical School|Faculdade de Ciências Médicas, NMS|FCM, Universidade Nova de Lisboa, Rua Camara Pestana, 6, 1150-082 Lisboa, Portugal

